# Phosphorylation modulates liquid-liquid phase separation of the SARS-CoV-2 N protein

**DOI:** 10.1101/2020.06.28.176248

**Authors:** Christopher R. Carlson, Jonathan B. Asfaha, Chloe M. Ghent, Conor J. Howard, Nairi Hartooni, David O. Morgan

## Abstract

The nucleocapsid (N) protein of coronaviruses serves two major functions: compaction of the RNA genome in the virion and regulation of viral gene transcription in the infected cell^1–3^. The N protein contains two globular RNA-binding domains surrounded by regions of intrinsic disorder^4^. Phosphorylation of the central disordered region is required for normal viral genome transcription^5,6^, which occurs in a cytoplasmic structure called the replication transcription complex (RTC)^7–11^. It is not known how phosphorylation controls N protein function. Here we show that the N protein of SARS-CoV-2, together with viral RNA, forms biomolecular condensates^12–15^. Unmodified N protein forms partially ordered gel-like structures that depend on multivalent RNA-protein and protein-protein interactions. Phosphorylation reduces a subset of these interactions, generating a more liquid-like droplet. We speculate that distinct oligomeric states support the two functions of the N protein: unmodified protein forms a structured oligomer that is suited for nucleocapsid assembly, and phosphorylated protein forms a liquid-like compartment for viral genome processing. Inhibitors of N protein phosphorylation could therefore serve as antiviral therapy.

## Introduction

Coronaviruses are enveloped viruses with a ~30 kb positive-sense single-stranded RNA genome, packed tightly inside the ~100 nm virion in a poorly-defined structure called the nucleocapsid^16–19^. Following viral entry and disassembly of the nucleocapsid, the genome is translated to produce RNA-dependent RNA polymerase and numerous non-structural proteins (Nsps)^1,20–22^.These proteins rearrange membranes of the endoplasmic reticulum to form the RTC^7–11^, which is thought to provide a scaffold for the viral proteins that perform genome replication and transcription, and which might also shield these processes from the host cell’s innate immune response.

Using genomic (+) RNA as a template, the viral polymerase produces (−) RNA transcripts of subgenomic regions encoding the four major viral structural proteins (S, E, M, and N)^1,20–22^. Subgenomic transcription involves a template-switching mechanism in which the polymerase completes transcription of a structural protein gene and then skips to a transcription-regulating sequence (TRS) at the 5’ end of the genome, resulting in subgenomic (−) RNA fragments – which are then transcribed to produce (+) RNA for translation. N protein is encoded by the most abundant subgenomic RNA and is translated at high levels early in infection. The N protein is among the most abundant viral proteins in the infected cell^2,3^ and accumulates in dynamic clusters at RTCs^11,23–25^, where it is thought to help promote the RNA structural rearrangements required for subgenomic transcription^5,26,27^.

The structural features of the N protein are well conserved among coronaviruses. The ~46 kDa N proteins of SARS-CoV and SARS-CoV-2 are ~90% identical. The N protein contains two globular domains, the N- and C-terminal domains (NTD and CTD), surrounded by intrinsically disordered regions^4^ (Figure 1a, Extended Data Fig. 1). N protein is highly basic (pI~10), and multiple RNA-binding sites are found throughout the protein^28^. The NTD is an RNA-binding domain^29–32^. The CTD forms a tightly-linked dimer with a large RNA-binding groove^33–36^, and the fundamental unit of N protein structure is a dimer^37,38^. Under some conditions, the dimer self-associates to form oligomers that depend on multiple protein regions^33,35,39–44^. The biochemical features and function of these oligomers are not known.

**Figure 1.**
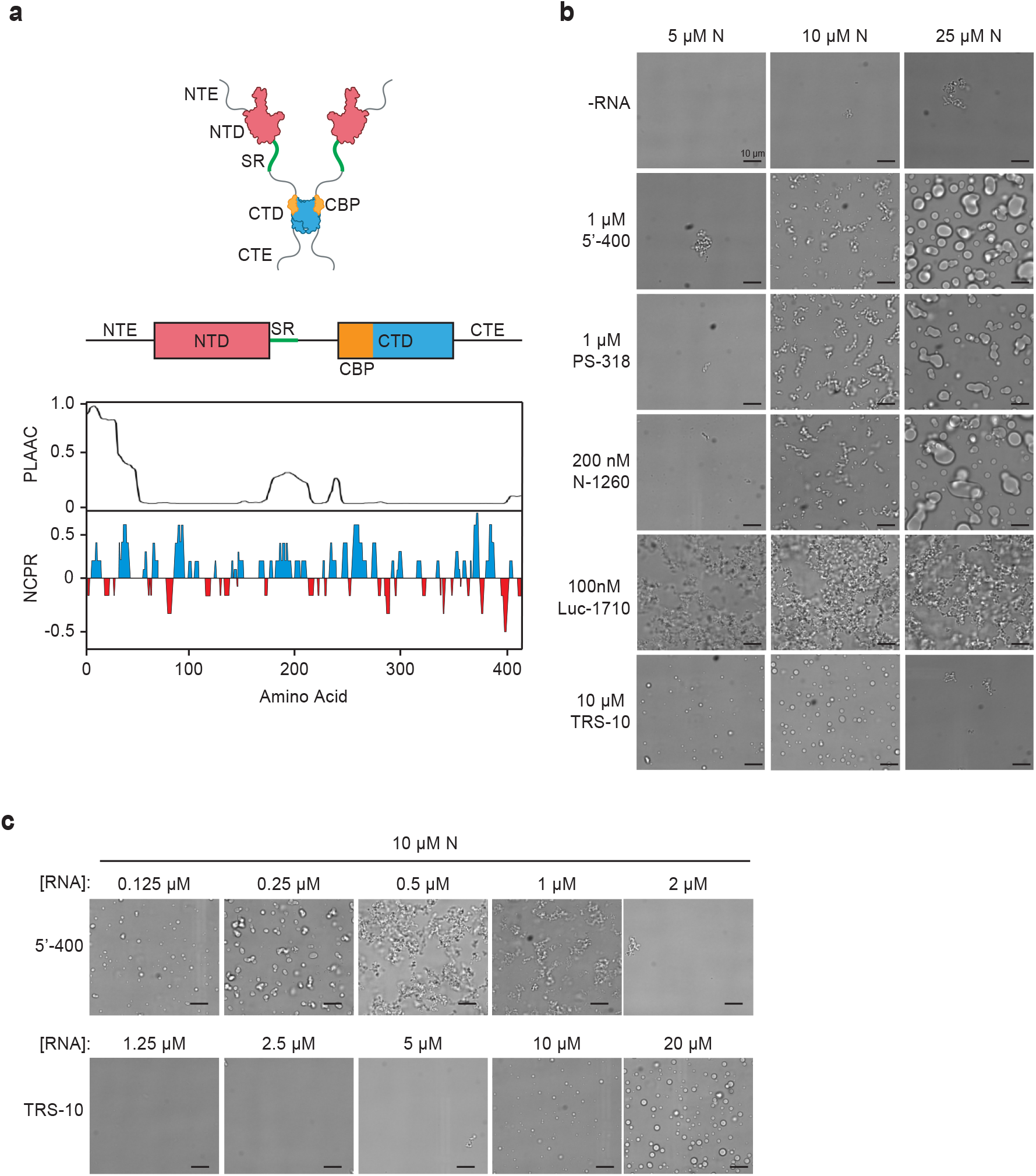
SARS-CoV-2 N protein forms biomolecular condensates in the presence of RNA. **a,** Top, schematic of N protein domain architecture. NTE, N-terminal extension; NTD, N-terminal domain; SR, SR region; CTD, C-terminal domain; CTE, C-terminal extension; CBP, CTD basic patch. Bottom, features of amino acid sequence. PLAAC, prion-like amino acid composition^78^, NCPR, net charge per residue. See Extended Data Fig. 1 for sequence. **b,** Light microscopy images of N protein condensates after 30 min incubation at room temperature with various RNA molecules. **c,** Condensate formation by N protein (10 μM) over a range of 5’-400 and TRS-10 RNA concentrations. All images are representative of multiple independent experiments; scale bar, 10 μm (**b, c**).

The central disordered linker contains a conserved serine-arginine (SR)-rich sequence that is likely to serve as a key regulatory hub in N protein function. Early in infection, the SR region is rapidly phosphorylated at multiple sites by cytoplasmic kinases^5,6,45–50^. Phosphorylation leads to association with the RNA helicase DDX1, which promotes RNA structural changes required for transcription of long subgenomic RNAs in the RTC^5^. Later in infection, nucleocapsid formation and viral assembly do not seem to require N protein phosphorylation, which is substantially reduced in the nucleocapsid of MHV and SARS-CoV virions^5,6^. We have little understanding of the molecular mechanisms by which phosphorylation influences N protein function.

## N protein of SARS-CoV-2 forms RNA-dependent biomolecular condensates

To gain a better understanding of N protein structure and function, we explored the oligomerization of the N protein from SARS-CoV-2, the causative agent of the ongoing COVID-19 pandemic. Purified N protein produced in bacteria migrated on gel filtration as a dimer in high salt but as a large oligomer in physiological salt (Extended Data Fig. 2a). Light microscopy revealed the presence of liquid-like droplets in the presence and absence of added RNA (Extended Data Fig. 2b). We noted, however, that purified N protein contained nucleic acid, even after RNAse treatment, raising the possibility that structures seen in the absence of added RNA were due to tightly-bound contaminating RNA. Following removal of RNA by protein denaturation and renaturation, the protein displayed few microscopic structures, but addition of viral RNA greatly enhanced the formation of structures similar to those in the native preparation (Fig. 1b). We conclude that RNA is required for the formation of the higher-order oligomers seen in the microscope. All subsequent studies were performed with denatured and renatured proteins (Extended Data Fig. 3a).

We first analyzed the effects of three mid-sized viral RNA fragments: (1) 5’-400, containing the 400 nt at the 5’ end of the SARS-CoV-2 genome, which is thought to include multiple secondary structure elements and the leader TRS^51–53^; (2) PS-318, a sequence near the end of ORF1b in SARS-CoV, proposed as a packaging sequence^54,55^ but of unknown function^51,56^; and (3) N-1260, containing the open reading frame of the N gene near the 3’ end of the SARS-CoV-2 genome. RNA encoding firefly luciferase (Luc-1710) was a nonviral control. N protein structures were analyzed by microscopy 30 minutes after addition of RNA at room temperature (Fig. 1b). At 10 μM N protein, the three viral RNAs rapidly generated branched networks of liquid-like beads. Higher N protein concentrations produced large liquid-like droplets several microns in diameter. Nonviral RNA led to amorphous filamentous aggregates with partial liquid-like appearance.

Incubation at higher temperature (30°C or 37°C) had little effect on droplet formation in a 30-minute incubation (Extended Data Fig. 3b), and longer incubations did not transform filamentous networks into droplets (Extended Data Fig. 3c). High salt dissolved N protein structures, indicating that they depend primarily on electrostatic interactions (Extended Data Fig. 3d).

N protein structures displayed different features at different ratios of N protein to 5’-400 RNA (Fig. 1c). At low RNA concentration (0.125 μM), 10 μM N protein formed small spherical droplets. Higher RNA concentrations led to filamentous structures. At RNA concentrations approaching that of N protein, no structures were formed. These results suggest that condensates depend on the crosslinking of multiple N proteins by a single RNA.

We tested the importance of multivalent RNA binding by measuring the effects of a 10-nucleotide RNA carrying the TRS sequence of SARS-CoV-2. The TRS sequence is thought to bind primarily to the NTD, with some contribution from the adjacent SR region^57,58^. Surprisingly, addition of the TRS RNA triggered the rapid formation of droplets, without the filamentous structures seen with longer viral RNAs (Fig. 1b, c). Also in contrast to results with longer RNAs, droplet formation was greatly reduced when the N protein was in molar excess over the TRS RNA (Fig. 1b, c), indicating that RNA-free N protein exerts a dominant inhibitory effect on the formation of TRS-bound oligomers. Droplets formed when TRS RNA was equimolar with or in excess of N protein (Fig. 1c), suggesting that these droplets depend on binding of a single TRS RNA to each N protein. Monovalent TRS binding appears to alter N protein structure to enhance low-affinity protein-protein interactions leading to droplet formation. In the more physiologically relevant context of long RNAs, these weak protein-protein interactions are presumably augmented by multivalent RNA-protein interactions.

We next explored the roles of N protein disordered regions. We analyzed mutant proteins lacking the following regions (Fig. 2a; Extended Data Fig. 1): (1) the 44-aa N-terminal extension (NTE), a poorly conserved prion-like sequence with a basic cluster that contributes to RNA binding^28^; (2) the 31-aa SR region, a basic segment implicated in RNA binding, oligomerization^44^, and phosphorylation^5,6,45–50^; (3) the 55-aa C-terminal extension (CTE), implicated in oligomerization^35,42–44^; and (4) the CTD basic patch (CBP), a 33-aa basic region that forms the RNA-binding groove on the surface of the CTD^33–35^, which can be deleted without affecting CTD dimer structure^36^.

**Figure 2.**
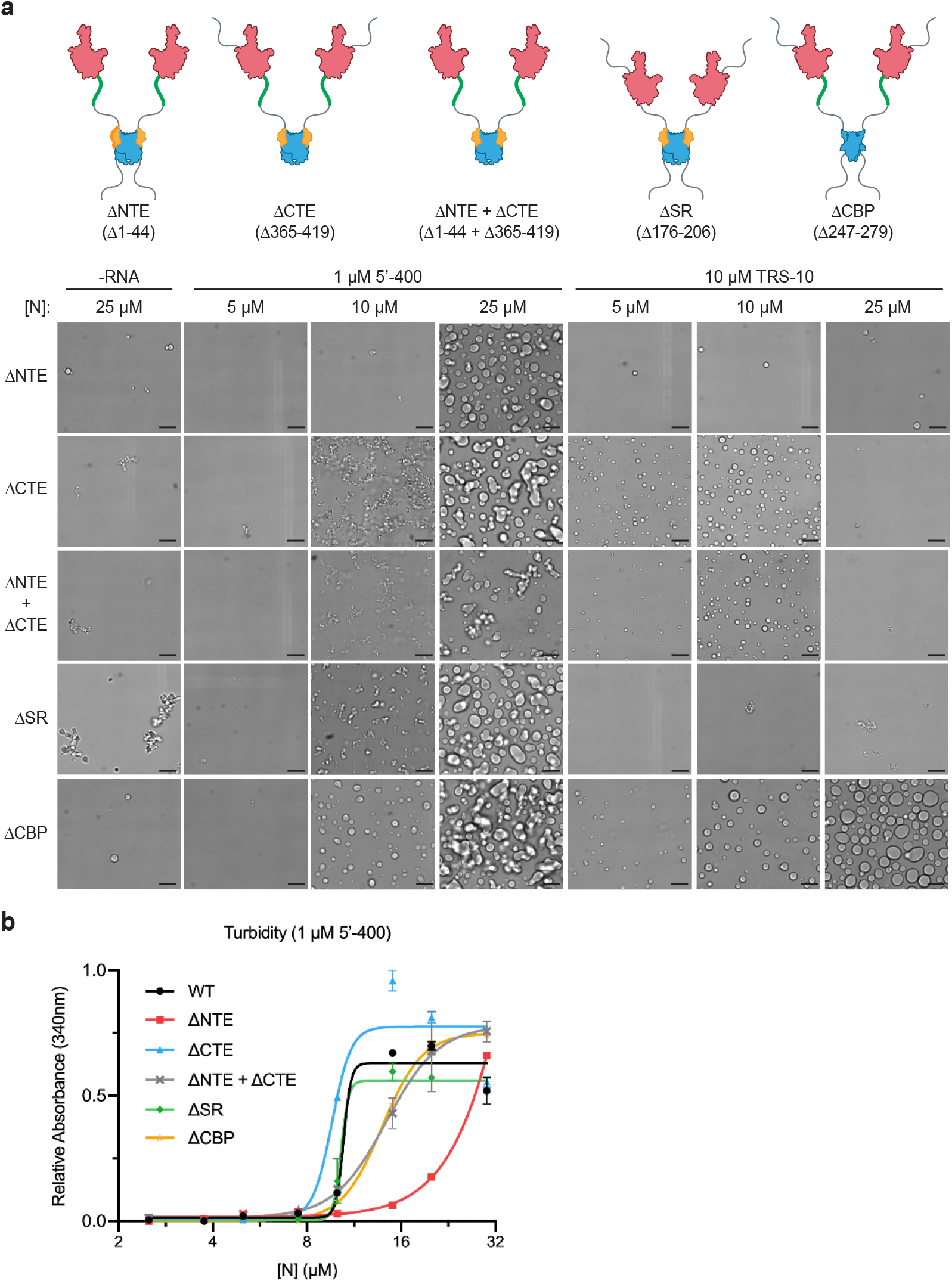
Disordered regions modulate N protein condensate formation. **a,** Top, schematics of N protein deletion mutants. Bottom, N protein condensates observed in the presence of 5’-400 RNA or TRS-10 RNA after a 30 min incubation at room temperature. Images are representative of multiple independent experiments. Scale bar, 10 μm. **b,** Absorbance at 340 nm was used to quantify the turbidity of N protein mixtures after a 15 min incubation at room temperature with 1μM 5’-400. Data points indicate mean +/− s.e.m. of duplicates; representative of two independent experiments.

When combined with viral 5’-400 RNA, none of the deletions completely prevented droplet formation at high protein concentrations (Fig. 2a), indicating that no single disordered segment is essential for the interactions that mediate droplet formation. CTE deletion stimulated the formation of abundant filaments, suggesting that this region normally inhibits interactions. Deletion of the NTE or CBP abolished filaments. Droplets were also observed after deletion of both the NTE and CTE, showing that the central regions are sufficient for droplet formation. Turbidity analyses (Fig. 2b) showed that full-length N protein structures increased abruptly between 5 and 10 μM, supporting a cooperative mechanism of oligomer assembly. CTE deletion reduced the saturating concentration and NTE deletion increased it, further supporting the negative and positive roles, respectively, of these regions. Deletion of both NTE and CTE resulted in an intermediate phenotype (Fig. 2b).

Further insights arose in studies of deletion mutants and the 10-nt TRS RNA (Fig. 2a). By minimizing the contribution of multivalent RNA binding, these studies illuminated critical protein-protein interactions that contribute to condensate formation. As in the experiments with long RNA, CTE deletion enhanced droplet formation, NTE deletion abolished it, and the double deletion had little effect, pointing to these regions as opposing but nonessential modulators of protein-protein interactions. As in the wild-type protein (Fig. 1b), a molar excess of N protein suppressed droplet formation by TRS RNA in most mutants; only the CBP deletion caused abundant droplets when the protein was in excess of RNA, suggesting that the CBP is responsible for the reduced droplet formation seen with high N protein concentrations. In contrast to results with long RNA, deletion of the SR region abolished TRS-mediated condensates (Fig. 2a). TRS binding to the NTD is known to be enhanced by the basic SR region, but SR deletion has only a moderate impact on affinity^57^. At the RNA concentration used in our experiments, it is unlikely that SR deletion abolished RNA binding. We therefore suspect that the SR region, perhaps in association with part of the RNA, is required for TRS-dependent droplet formation because it mediates a weak interaction with another N protein^44^.

## Phosphorylation promotes more liquid-like N protein condensates

N protein phosphorylation depends on a poorly understood collaboration between multiple kinases^5,6,49^. A central player is the abundant cytoplasmic kinase GSK-3, which generally phosphorylates serines or threonines four residues upstream of pre-phosphorylated ‘priming’ sites^59^. Studies of the N protein of SARS-CoV^6^ support the presence of two priming sites, P1 and P2 (Fig. 3a), where phosphorylation initiates a series of GSK-3-mediated phosphorylation events, each primed by the previous site, resulting in a high density of up to ten phosphates. The kinases responsible for priming phosphorylation are not known, but the P2 site (S206 in SARS-CoV-2) is a strong consensus sequence (S/T-P-x-K/R) for Cdk1, a major cell cycle kinase^49^.

**Figure 3.**
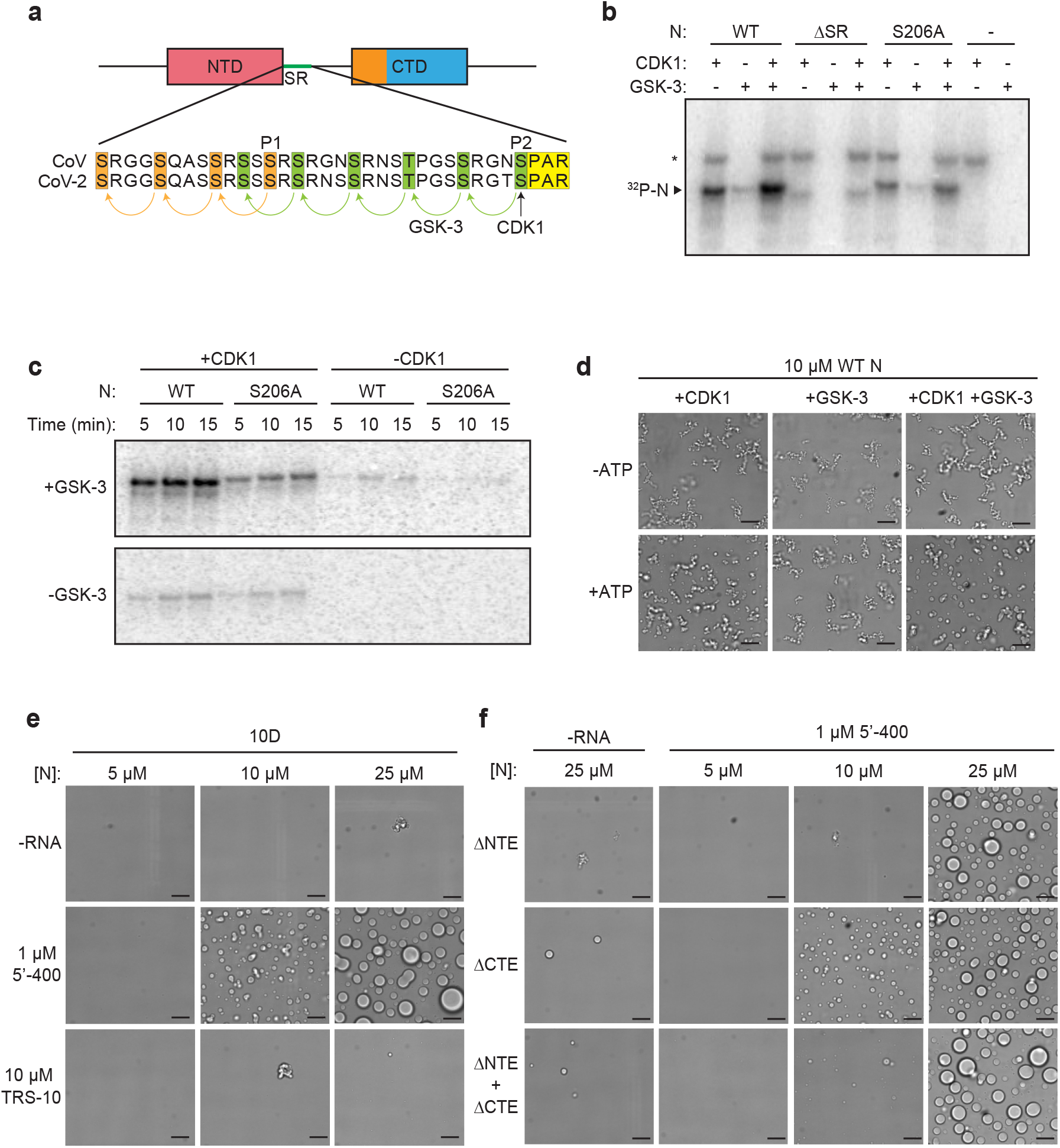
Phosphorylation modulates N protein condensate properties. **a,** Sequences of the SR regions of SARS-CoV N protein (aa 177-210) and SARS-CoV-2 N protein (aa 176-209). Proposed priming sites (P1 and P2) for GSK-3 are indicated^6^. P2 (S206 in CoV-2) is a Cdk consensus site (yellow), where phosphorylation is thought to prime sequential phosphorylation (arrows) of 5 upstream sites (green) by GSK-3. P1 phosphorylation by an unknown kinase primes phosphorylation at 3 upstream sites (orange). **b,** The indicated N protein variants were incubated 30 min with Cdk1-cyclin B1 and/or GSK-3 and radiolabeled ATP, and reaction products were analyzed by SDS-PAGE and autoradiography. Radiolabeled N protein is indicated. Asterisk indicates cyclin B1 autophosphorylation. **c,** N protein was incubated overnight with unlabeled ATP and Cdk1-cyclin B1 (left lanes) or no kinase (right lanes), desalted, and incubated with or without GSK-3 and radiolabeled ATP. Reaction products were analyzed by SDS-PAGE and autoradiography. **d,** 10 μM N protein was incubated 2 h with Cdk1-cyclin B1 and GSK-3 in the presence or absence of ATP, dialyzed into droplet buffer overnight, and then mixed with 1 μM 5’-400 RNA. After 30 min, N protein condensates were analyzed by light microscopy. **e,** Images of N protein 10D mutant following 30 min incubation with or without 1 μM 5’-400 or 10 μM TRS-10 RNA. **f,** Images of 10D mutants with the indicated deletions, incubated with or without 1 μM 5’-400 RNA. All results are representative of multiple independent experiments; scale bar, 10 μm (**d-f**).

To produce phosphorylated N protein, we first tested the possibility that Cdk1 primes the protein for subsequent phosphorylation by GSK-3. We found that Cdk1-cyclin B1 phosphorylated N protein in the SR region, and mutation of S206 reduced Cdk1-dependent phosphorylation (Fig. 3b). Phosphorylation might also occur at T198, a nearby Cdk1 consensus site. A combination of Cdk1 and GSK-3 enhanced phosphorylation. Clear evidence for priming by Cdk1 was obtained by extensive unlabeled phosphorylation by Cdk1, followed by analysis of radiolabeled phosphorylation with GSK-3 (Fig. 3c).

Phosphorylation of N protein with a combination of Cdk1 and GSK-3 reduced filamentous structures and promoted the formation of liquid-like droplets (Fig. 3d). GSK-3 alone had no effect, whereas Cdk1 alone promoted droplets to a small extent. We conclude that phosphorylation in the SR region shifts the behavior of N protein to promote the formation of liquid-like condensates.

We explored the role of phosphorylation in depth with studies of a phosphomimetic mutant in which the ten serines and threonines in the SR region were replaced with aspartate (the 10D mutant). When combined with the 5’-400 viral RNA, the 10D protein rapidly formed condensates with a spherical droplet morphology that was clearly distinct from the filamentous structures of the wild-type protein (Fig. 3e). All three large viral RNAs were effective in driving droplet formation, although N-1260 appeared to reduce the saturating concentration (Extended Data Fig. 4a). NTE deletion in the 10D protein reduced droplet formation, showing once again the positive role of this region (Fig. 3f). Importantly, the 10D mutation had a greater impact on condensate morphology than an SR deletion (Fig. 2a), suggesting that phosphorylation does not just block SR function but might also interfere with other interactions. One possibility, for example, is that the abundant negative charge of the phosphorylated SR region interacts intramolecularly with one or more of the positively-charged patches on the adjacent NTD or CTD, thereby interfering with multivalent RNA-protein interactions.

Droplet formation by TRS RNA was abolished in the 10D mutant (Fig. 3e), just as we observed with TRS RNA and the SR deletion (Fig. 2a). These results support the notion that phosphorylation blocks the weak protein-protein interactions mediated by the SR region.

Phosphorylated N protein is thought to be localized at the RTC^48^. The N protein of mouse hepatitis virus (MHV) is known to interact directly with the N-terminal Ubl1 domain of Nsp3^23,60^, a large transmembrane protein localized to RTC membranes^23,60^. We found that a GFP-tagged Ubl1 domain of SARS-CoV-2 Nsp3 partitioned into N protein droplets and filamentous structures (Extended Data Fig. 4b), providing a potential mechanism for association of the RTC with N protein condensates.

To gain a better understanding of the properties of N protein structures, we analyzed the fusion dynamics of different structures over time. During a short (90 s) time course, the filamentous structures of 10 μM wild-type protein remained immobile and did not fuse, while the spherical droplets of the 10D mutant were highly dynamic and fused rapidly (Fig. 4a). We also found that the droplet-like structures at higher concentrations of unmodified protein displayed relatively slow fusion activity compared to the more dynamic droplets of phosphorylated N protein (Fig. 4b). Thus, unmodified protein forms gel-like condensates that are relatively rigid, while phosphorylated N protein forms condensates that behave more like liquid droplets.

**Figure 4.**
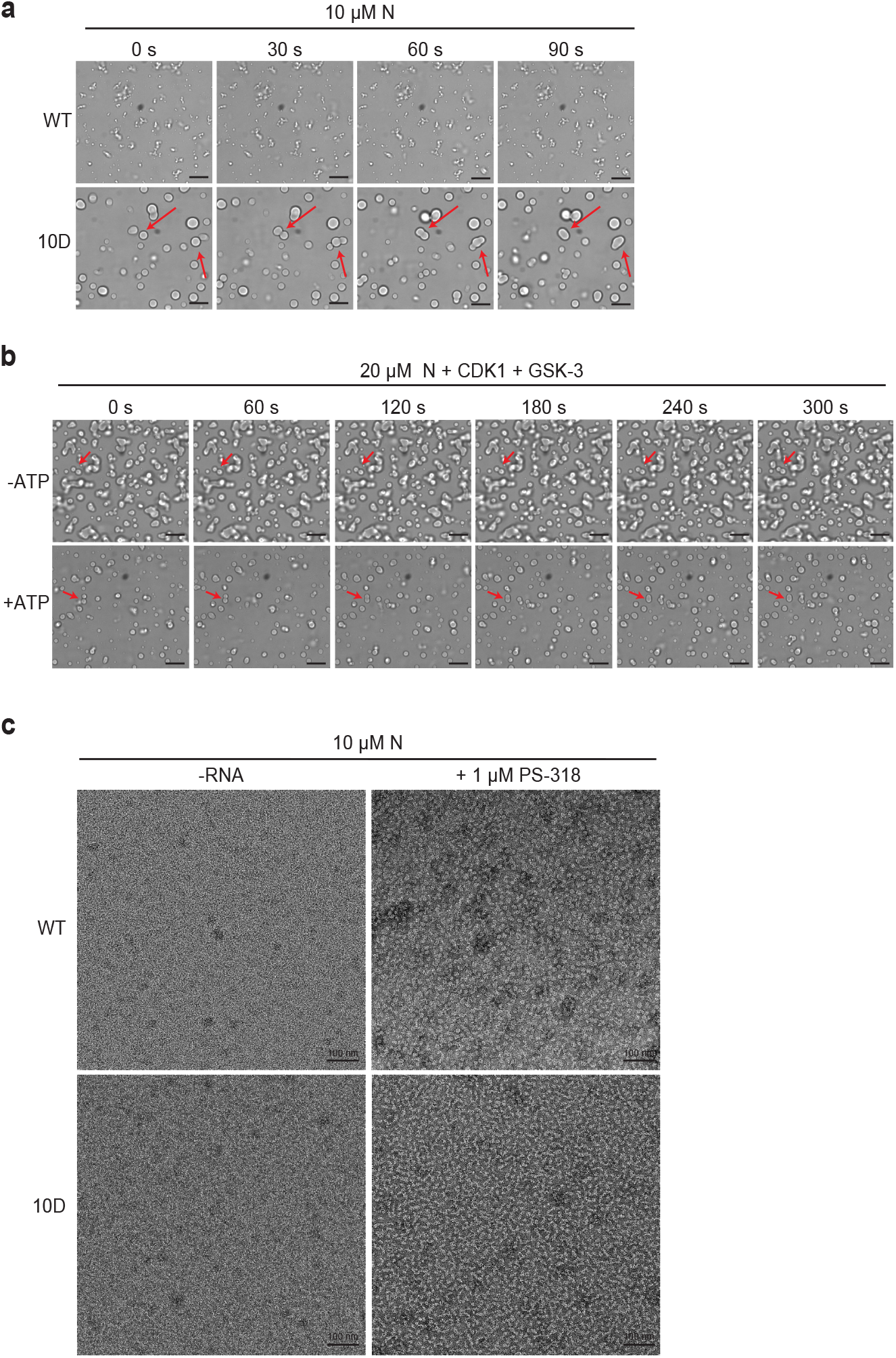
Phosphorylation of N protein promotes liquid-like behavior. **a,** N protein (10 μM wild-type [WT] or 10D) was mixed with 1 μM 5’-400 RNA for 20 min, and images were taken at 30 s intervals. Arrows indicate droplet fusion events in the 10D mutant. No fusion events were observed in WT structures. **b,** 20 μM N protein was phosphorylated with Cdk1-cyclin B1 and GSK-3 as in Fig. 3d, incubated with 1 μM 5’-400 RNA for 20 min, and imaged every 60 s. Arrows indicate droplet fusion events. Images are representative of multiple independent experiments; scale bar, 10 μm. **c,** 10 μM N protein (WT or 10D) was incubated without or with 1 μM PS-318 RNA for 15 min prior to analysis by negative-stain electron microscopy. Images are representative of three independent experiments. Scale bar, 100 nm.

We used negative-stain electron microscopy to further analyze mixtures of 10 μM N protein and 1 μM RNA (Fig. 4c). These images represent the saturated protein-RNA solution surrounding condensates. Unmodified wild-type protein and viral RNA formed uniform particles of ~20 nm diameter, reminiscent of structures seen in previous studies of partially disrupted MHV nucleocapsids^17^. 2D classification of these particles revealed a uniform size and architecture (Extended Data Fig. 4c). Thus, the gel-like filamentous condensates of the unmodified protein – and possibly the nucleocapsid – are likely to be assembled on a foundation of discrete structural building blocks. In contrast, the mixture of 10D mutant and RNA formed nonuniform, diffuse chains (Fig. 4c), suggesting that phosphorylation disrupts the structural units of the unmodified protein to help create a more liquid-like condensate.

## Discussion

We conclude that the N protein of SARS-CoV-2, together with viral RNA, assembles into multiple structural forms that depend on a complex blend of intra- and intermolecular interactions. The more rigid filamentous condensates of the unmodified protein likely depend on high-avidity interactions mediated by multivalent RNA-protein and protein-protein interactions. The latter might include prion-like interactions between NTEs, binding of the SR region to the CTD^44^, or helical CTD polymers that depend on the CBP^17,33^. Long RNAs augment these protein-protein interactions by interacting with the numerous RNA-binding sites on the protein. Our results also support a model in which phosphorylation of the SR region blocks SR-mediated protein-protein interactions and interferes intramolecularly with RNA binding at other sites, resulting in a loss of affinity that generates a more dynamic liquid droplet.

The different forms of N protein oligomers seem well suited for its two major functions. In the nucleocapsid, where extremely compact RNA packaging is the goal, the organized structures of the unmodified protein could represent an underlying structural framework that is supplemented by liquid-like condensation – much like chromosome packaging depends on underlying nucleosome structure and the liquid-like behavior of chromatin proteins^61,62^.

The more liquid-like behavior of phosphorylated N protein droplets might be particularly important at the RTC. There is abundant evidence linking N protein phosphorylation with localization and function in the RTC. N protein is fully phosphorylated soon after synthesis and rapidly associates with large membrane structures that presumably represent the RTC^48^. N protein is the only viral structural protein that localizes to the RTC^63^, where it is seen in immunoelectron microscopy adjacent to the double-membrane vesicles and convoluted membranes of this organelle^11^. GFP-tagged N protein forms large clusters at RTCs, and fluorescence recovery after photobleaching indicates that N protein is dynamically associated with these clusters^25^. N protein helps control subgenomic transcription in the RTC, and inhibition of phosphorylation blocks this function^5,26,27^. With these lines of evidence in mind, our work points to the possibility that a liquid-like matrix of phosphorylated N protein and loosely bound RNA, linked to RTC membranes by Nsp3, provides a compartment to concentrate and protect the viral replication and transcription machinery. Similar mechanisms are likely to exist in negative-sense RNA viruses, where replication is focused in dynamic biomolecular condensates 64-66. In Measles virus, these condensates have also been implicated in nucleocapsid assembly^67^. We also note that others have recently observed SARS-CoV-2 N protein condensates^68–71^.

Small chemical inhibitors of GSK-3 kinase activity disrupt MHV genome processing and reduce the production of virions by MHV- or SARS-CoV-infected cells^5,6^, consistent with the importance of N protein phosphorylation in genome replication. These inhibitors, perhaps together with inhibitors of priming kinases such as Cdk1, have the potential to serve as antiviral therapy in the early stages of COVID-19.

## Methods

### Plasmid construction and RNA preparation

All expression vectors were constructed by Gibson assembly and confirmed by sequencing. Codon-optimized DNAs encoding SARS-CoV-2 N protein (aa 1-419) and Nsp3 Ubl1 domain (aa 819-920 of polyprotein 1a) were ordered as gBlocks from Integrated DNA Technologies (IDT) and cloned into pET28a expression vectors. N protein mutants were constructed by site-directed mutagenesis and Gibson assembly. A 6xHis-SUMO tag was added to the N-terminus of all N protein constructs. Nsp3 Ubl1 domain was engineered with a C-terminal GFP-2xStrep tag. Codon-optimized DNAs encoding human Cdk1 (untagged) and Cyclin B1 (N-terminal 6xHis-4xMyc tag) were ordered as gBlocks from IDT and cloned individually into the baculovirus expression vector pLIB for construction of recombinant baculoviruses^72^. Human GSK-3β was purchased from Promega (V1991).

Sequences of the three viral RNAs and firefly luciferase RNA are shown in Extended Data Fig. 5. Templates for transcription in vitro of 5’-400 RNA and PS-318 RNA were ordered as gBlocks from IDT and PCR-amplified with a 5’ primer carrying a T7 promoter sequence. The template for N-1260 was PCR-amplified with a 5’ primer containing a T7 promoter from a PET28a vector carrying PCR-amplified ORF DNA from the 2019-nCoV_N positive control plasmid from IDT. The N-1260 RNA includes 152 nucleotides from the pET28a plasmid backbone. All RNA synthesis was performed using the HiScribe T7 High Yield RNA synthesis kit (NEB E2040S) according to the manufacture’s protocol. Luc-1710 RNA was included as a positive control template in the RNA synthesis kit. The SARS-CoV-2 TRS RNA (UCUAAACGAA, tagged with FAM fluorescent dye) was ordered from IDT.

### Protein purification

N protein vectors were transformed into *E. coli* BL21 star (Thermo #C601003) for expression. Freshly transformed cells were grown in TB-Kanamycin (50 μg/ml) to OD 0.4 at 37°C. The temperature was lowered to 16°C until the cells reached a density of 0.8. Protein expression was then induced with 0.4 mM IPTG for 16 h. Harvested cells were washed with PBS, snap frozen in LN_2_, and stored at −80°C until lysis.

To remove contaminating nucleic acid, N protein was purified under denaturing conditions as previously described^47,73^. Frozen cell pellets were thawed and resuspended in buffer A (50 mM Hepes pH 7.5, 500 mM NaCl, 10% glycerol, 20 mM imidazole, 6 M urea) and lysed by sonication on ice. Lysed cells were centrifuged at 30,000 rpm for 30 min at 4°C to remove cell debris. Clarified lysate was added to Ni-NTA agarose beads (Qiagen) and incubated for 45 min at 4°C. Ni-NTA beads were then washed with 3 × 10 bed volumes of buffer A, and N protein was eluted with 3 bed volumes buffer B (50 mM Hepes pH 7.5, 500 mM NaCl, 10% glycerol, 250 mM imidazole, 6 M urea). The eluate was concentrated to ~1 ml using centrifugal concentrators (30 kDa cutoff, Amicon) and renatured by overnight dialysis against 2 l buffer C (50 mM Hepes pH 7.5, 500 mM NaCl, 10% glycerol) at 4°C. 100 μg recombinant Ulp1 catalytic domain (expressed in and purified from bacteria) was added to renatured protein for 10 min on ice to cleave the 6xHis-SUMO tag. Mutants lacking the NTE were not cleaved as efficiently by Ulp1, and complete cleavage required incubation with Ulp1 at 25°C for 4 h. Cleaved protein was then centrifuged at 15,000 rpm for 10 min and injected onto a Superdex 200 10/300 size exclusion column equilibrated in buffer C. Peak fractions were pooled, frozen in LN_2_, and stored at −80°C.

In early experiments, N protein was purified under native conditions. Frozen cell pellets were thawed and resuspended in buffer D (50 mM Hepes pH 7.5, 500 mM NaCl, 10% glycerol, 20 mM imidazole) supplemented with benzonase, cOmplete EDTA-free protease inhibitor cocktail (Roche), and 1 mM phenylmethylsulfonyl fluoride (PMSF). Cells were lysed by sonication on ice and centrifuged at 30,000 rpm for 30 min at 4°C. Clarified lysate was added to Ni-NTA agarose beads and incubated for 45 min at 4°C. Ni-NTA beads were then washed with 3 × 10 bed volumes buffer D, and N protein was eluted with 3 bed volumes buffer D + 500 mM imidazole.

The eluate was concentrated to ~1 ml using centrifugal concentrators (30 kDa cutoff, Amicon) and injected onto a Superdex 200 10/300 size exclusion column. Peak fractions were pooled, snap frozen in LN_2_, and stored at −80°C. Protein concentration was measured by nanodrop, and a major A260 peak was observed. The A260 peak was insensitive to treatment with DNase I, RNase A, RNase H, or benzonase. Additionally, small RNA species were observed on native TBE gels stained with Sybr gold.

Nsp3 Ubl1-GFP was expressed in *E. coli* as described above. Frozen cell pellets were thawed and resuspended in buffer E (50 mM Hepes pH 7.5, 150 mM NaCl, 10% glycerol, 1 mM DTT) supplemented with benzonase, cOmplete EDTA-free protease inhibitor cocktail (Roche), and 1 mM PMSF. Cells were lysed by sonication on ice and centrifuged at 30,000 rpm for 30 min at 4°C. Clarified lysate was then added to a 5 ml StrepTrap HP prepacked column (GE). The column was washed with 10 column volumes buffer E and eluted with 4 column volumes buffer E supplemented with 2.5 mM desthiobiotin. Peak fractions were pooled, snap frozen in LN_2_, and stored at −80°C.

Cdk1-cyclin B1 complexes were prepared as follows. Two 666 ml cultures of SF-900 cells (1.6×10^6^ cells/ml) were infected separately with Cdk1 or cyclin B1 baculovirus and harvested after 48 h. The two cell pellets were frozen in LN_2_. Frozen pellets were thawed and each resuspended in 20 ml lysis buffer (50 mM Hepes pH 7.5, 300 mM NaCl, 10% glycerol, 20 mM imidazole, benzonase, and cOmplete EDTA-free protease inhibitor cocktail). Cells were lysed by sonication. To generate an active Cdk1-cyclin B1 complex phosphorylated by Cdk-activating kinase in the lysate^74^, the two lysates were combined, brought to 5 mM ATP and 10 mM MgCl_2_, and incubated at room temperature for 20 min. The combined lysates were centrifuged at 55,000 rpm for 45 min at 4°C. The supernatant was filtered and passed over a HisTRAP nickel affinity column, washed with wash buffer (50 mM Hepes pH 7.5, 300 mM NaCl, 10% glycerol) and eluted with the same buffer plus 200 mM imidazole. Peak fractions were pooled, concentrated to 0.75 ml, and injected on an S200 size exclusion column in wash buffer. Peak fractions containing the Cdk1-cyclin B1 complex were pooled, concentrated to 1.5 mg/ml, and snap frozen in LN_2_.

### Light microscopy

Glass was prepared as described previously^75^. Individual wells in a 384-well glass bottom plate (Greiner #781892) were incubated with 2% Hellmanex detergent for 1 h. Wells were then washed 3 times with 100 μl ddH_2_O. 1 M NaOH was added to the glass for 30 min, followed by washing 3 times with 100 μl ddH_2_O. The glass was dried, and 20 mg/ml PEG silane dissolved in 95% EtOH was added to individual wells and incubated overnight (~16 h). The glass was then washed 3 times with 100 μl ddH_2_O and dried before sample addition.

The day prior to imaging, protein was thawed and dialyzed against droplet buffer (25 mM Hepes pH 7.5, 70 mM KCl) overnight at 4°C. Protein concentration was then quantified by nanodrop. Reactions containing protein and RNA were combined and mixed immediately before adding to individual wells in the PEG-treated 384-well plate. All reactions were incubated for 30 min at room temperature, unless otherwise indicated, before imaging on a Zeiss Axiovert 200M microscope with a 40x oil objective.

### Turbidity analysis

Protein was dialyzed into droplet buffer the night before performing turbidity measurements, as stated above. Protein was then serially diluted into room temperature droplet buffer. After dilution, RNA was added and mixtures were incubated at room temperature for 15 min before measuring the absorbance at 340 nm on a Spectramax M5 plate reader.

### Kinase reactions

1 μM N protein was incubated with either 0.14 μM GSK3-β, 0.91 μM Cdk1-Cyclin B1, or both, in a 20 μl reaction mixture containing 10 mM MgCl_2_, 1 mM DTT, 500 μM ATP, 25 mM Hepes pH 7.8, 50 mM NaCl, and 0.001 mCi/ml ^32^P-γ-ATP. After incubation at 30°C for 30 min, reactions were quenched with 10 μl SDS loading buffer for analysis by SDS-PAGE and visualization with a Phosphorimager. For the priming experiment in Fig. 3c, 19 μM of each N protein variant was incubated with 2.5 μM Cdk1-Cyclin B1 overnight at 25°C in kinase buffer A (20 mM Hepes pH 7.4, 300 mM NaCl, 20 mM MgCl_2_, 1 mM DTT, 1 mM ATP). Reactions were desalted into kinase buffer B (20 mM Hepes pH 7.4, 20 mM MgCl_2_, 1 mM DTT, 500 μM ATP). 10 μM of desalted N protein was incubated with 0.17 μM GSK3-β and 0.05 μCi ^32^P-γ-ATP. After various times, reactions were quenched with SDS loading buffer for analysis by SDS-PAGE and visualization with a Phosphorimager.

Phosphorylated protein for droplet analysis was prepared in reactions containing 25 μM N protein, 1 μM Cdk1-cyclin B1, 0.2 μM GSK-3β, 2 mM ATP, 20 mM MgCl_2_, 15 mM Hepes pH 7.5, 300 mM NaCl, 5% glycerol, 1 mM DTT, 8 mM phosphocreatine (Sigma 10621714001), and 0.016 mg/ml creatine kinase (Sigma C3755). The reaction was incubated for 2 h at 30°C and then dialyzed overnight into droplet buffer at 4°C. RNA addition and droplet visualization were carried out as described above.

### Electron microscopy

N protein (10 μM) and RNA (1 μM PS-318) were mixed in 10 μl droplet buffer and incubated 15 min at room temperature. 3.5 μl of this solution was adsorbed onto glow-discharged (PELCO EasiGlow, 15 mA, 0.39 mBar, 30 s) carbon-coated grids (200-400 mesh copper, Quantifoil) for 1 min at room temperature. Sample was blotted off, stained and blotted 5 times with 0.75% uranyl formate, and allowed to air dry. Negative stain images were collected with a Tecnai T12 microscope (FEI) with a LaB6 filament, operated at 120 kV, and a Gatan Ultrascan CCD camera (final pixel size 2.21 Å). Contrast Transfer Function (CTF) estimation was performed with CTFFIND4^76^. Automated particle picking, CTF-correction, and two-dimensional averaging and classification were performed in RELION3^77^.

## Data Availability

The data that support the findings of this study are available from the corresponding author upon request.

## Acknowledgements

We thank Geeta Narlikar, Sy Redding, Alan Frankel, and Adam Frost for discussions, and Madeline Keenen and Emily Wong for reagents and technical advice. This work was supported by the National Institute of General Medical Sciences (R35-GM118053) and the UCSF Program for Breakthrough Biomedical Research, which is partially funded by the Sandler Foundation.

## Author contributions

C.R.C., J.B.A., and C.M.G. contributed to conceptualization, experimental design, and generation of results; N.H. contributed to conceptualization and experimental design; C.J.H. performed EM analysis; D.O.M. provided guidance and wrote the paper with contributions from all authors.

**Extended Data Fig. 1.**
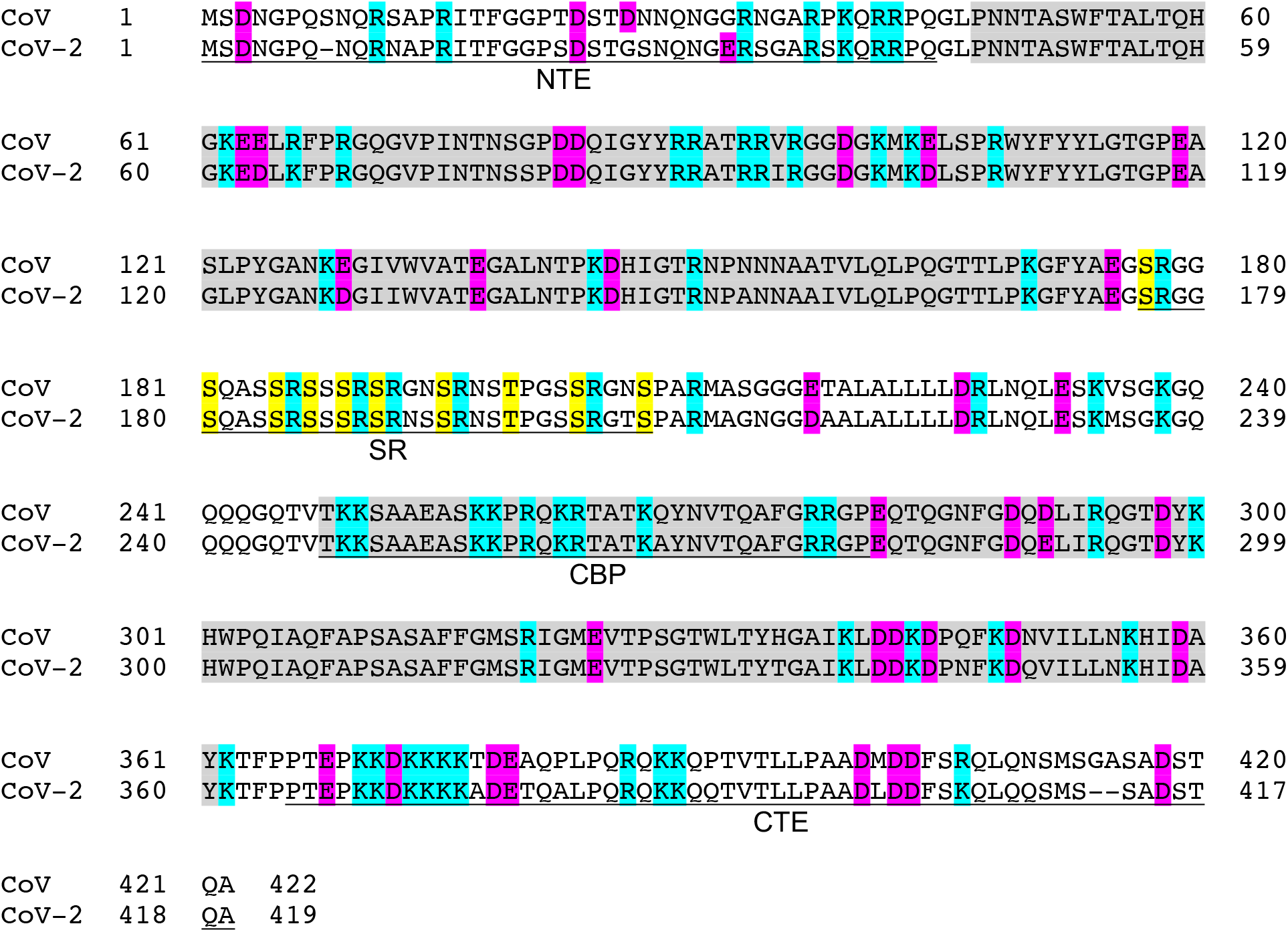
Amino acid sequences of N proteins from SARS-CoV and SARS-CoV-2. The two globular domains, NTD and CTD, are highlighted in gray. Underlining indicates the four regions analyzed by deletion mutants. Acidic residues highlighted in pink, basic residues highlighted in blue, and phosphorylation sites of the SR region highlighted in yellow.

**Extended Data Fig. 2.**
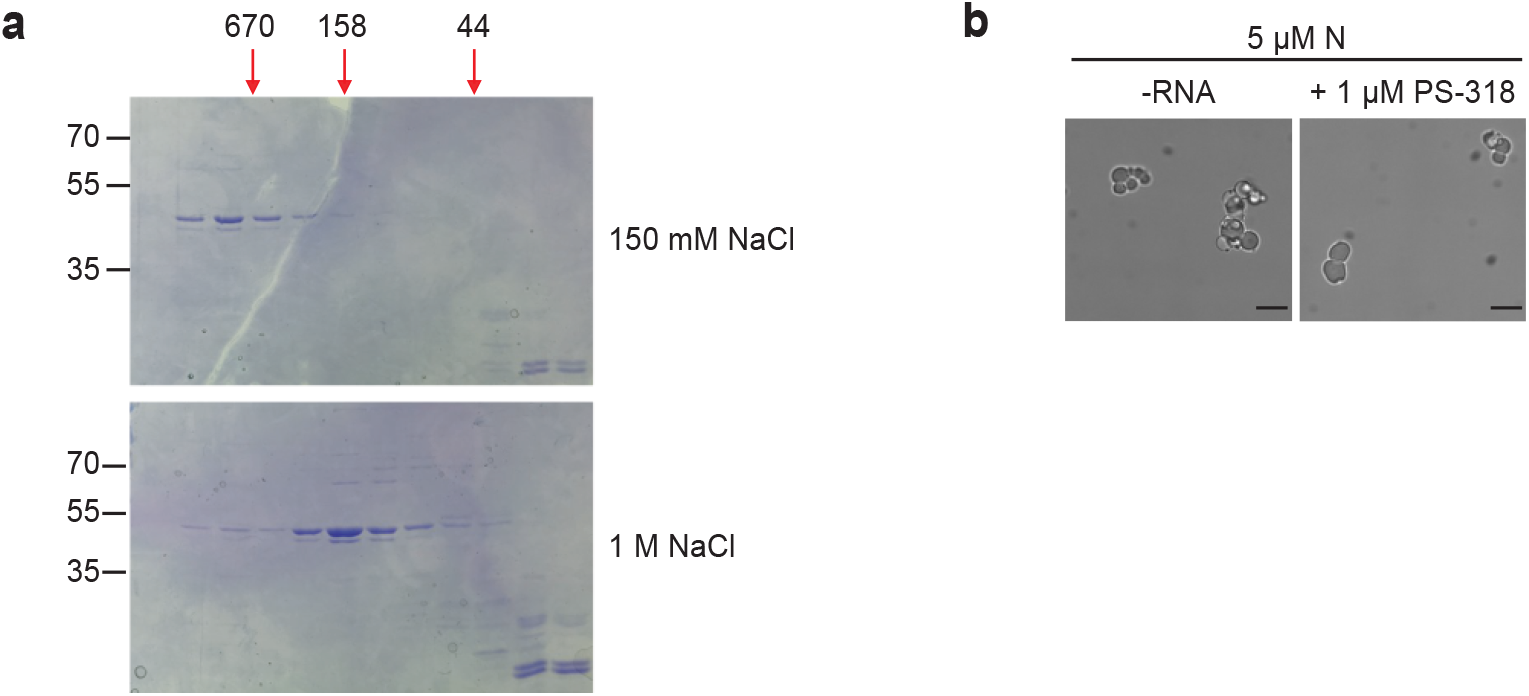
Native N protein preparations form condensates without added RNA. **a,** Superdex 200 gel filtration analysis of native N protein at 150 mM or 1 M NaCl, purified under non-denaturing conditions. Arrows at top indicate migration of molecular weight standards (kDa). Fractions were analyzed by SDS-PAGE and Coomassie Blue staining. **b,** Representative images of 5 μM native N protein after incubation in the presence or absence of 1 μM PS-318 RNA. Scale bar, 10 μm.

**Extended Data Fig. 3.**
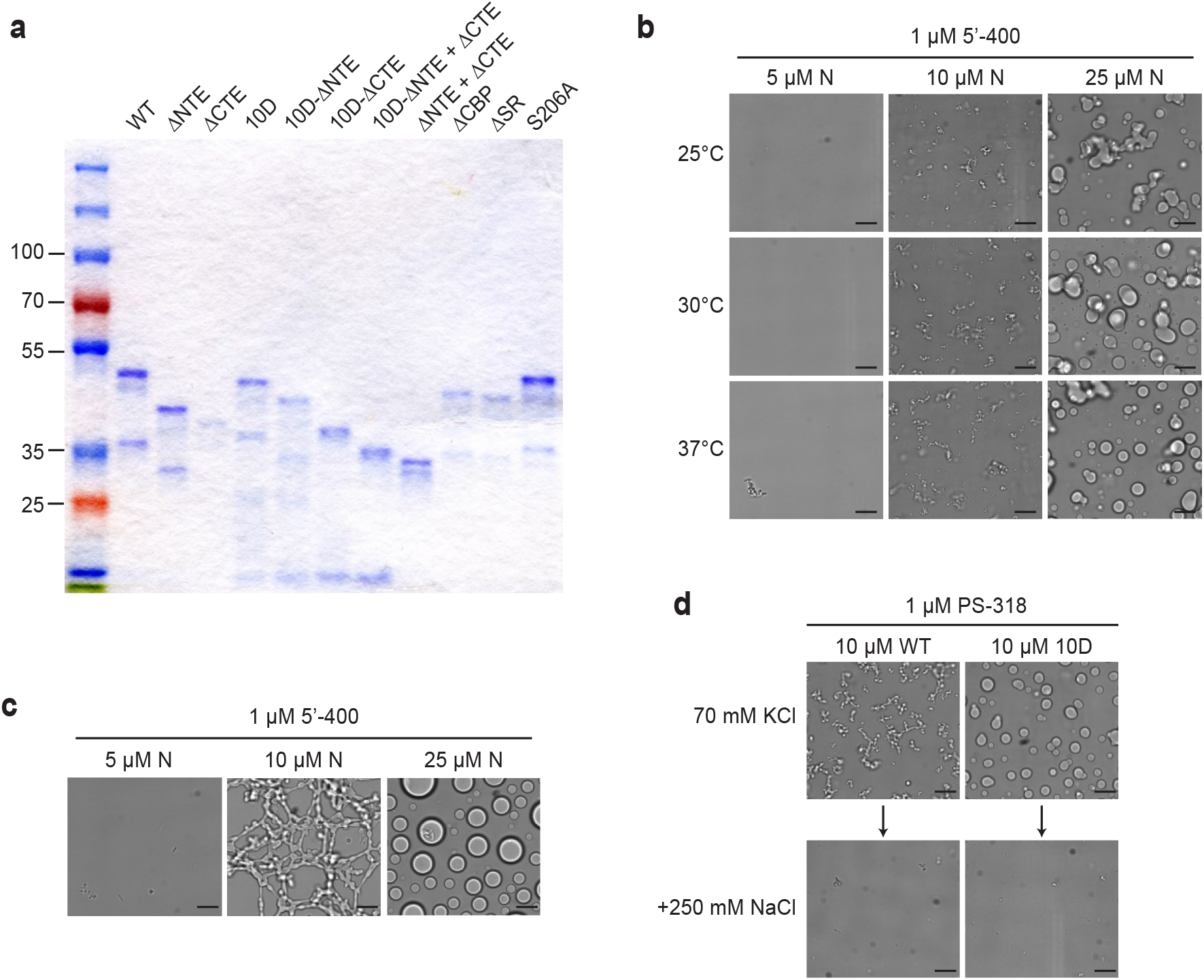
Characterization of N protein condensates. **a,** SDS-PAGE analysis of all N protein mutants used in this study, stained with Coomassie Blue. **b,** N protein was incubated at the indicated temperature for 30 min in the presence of 1 μM 5’-400 RNA. Scale bar, 10 μm. **c,** N protein was incubated with 1 μM 5’-400 for 16 h at room temperature. Scale bar, 10 μm. **d,** Condensates of 10 μM WT or 10D N protein were formed in droplet buffer (70 mM KCl) by incubation with 1 μM PS-318 RNA for 30 min and imaged. NaCl was then added to a final concentration of 250 mM for 15 min before imaging again.

**Extended Data Fig. 4.**
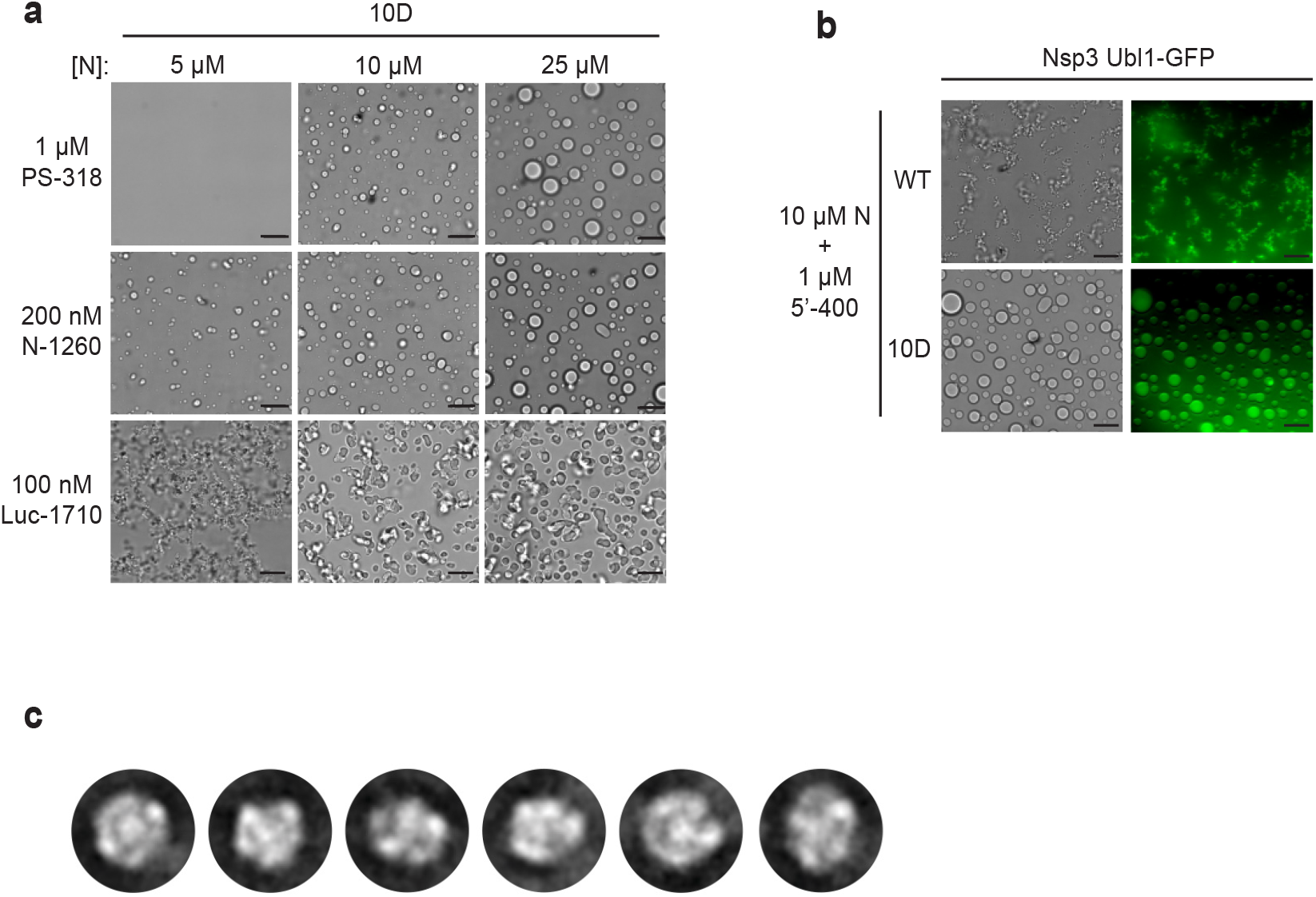
Characterization of N protein condensates. **a,** Images of N protein 10D mutant following 30 min incubation with the indicated RNAs. Scale bar, 10 μm. **b,** 10 μM wild-type (WT) or 10D N protein was incubated with 1 μM 5’-400 RNA for 10 min. Nsp3 Ubl1-GFP was then added to a concentration of 1 μM and incubated for an additional 15 min before imaging in brightfield (left) or fluorescence (right). **c,** 2D class averages of particles from the EM analysis of wild-type N protein and PS-318 RNA shown in Fig. 4c.

**Extended Data Fig. 5.**
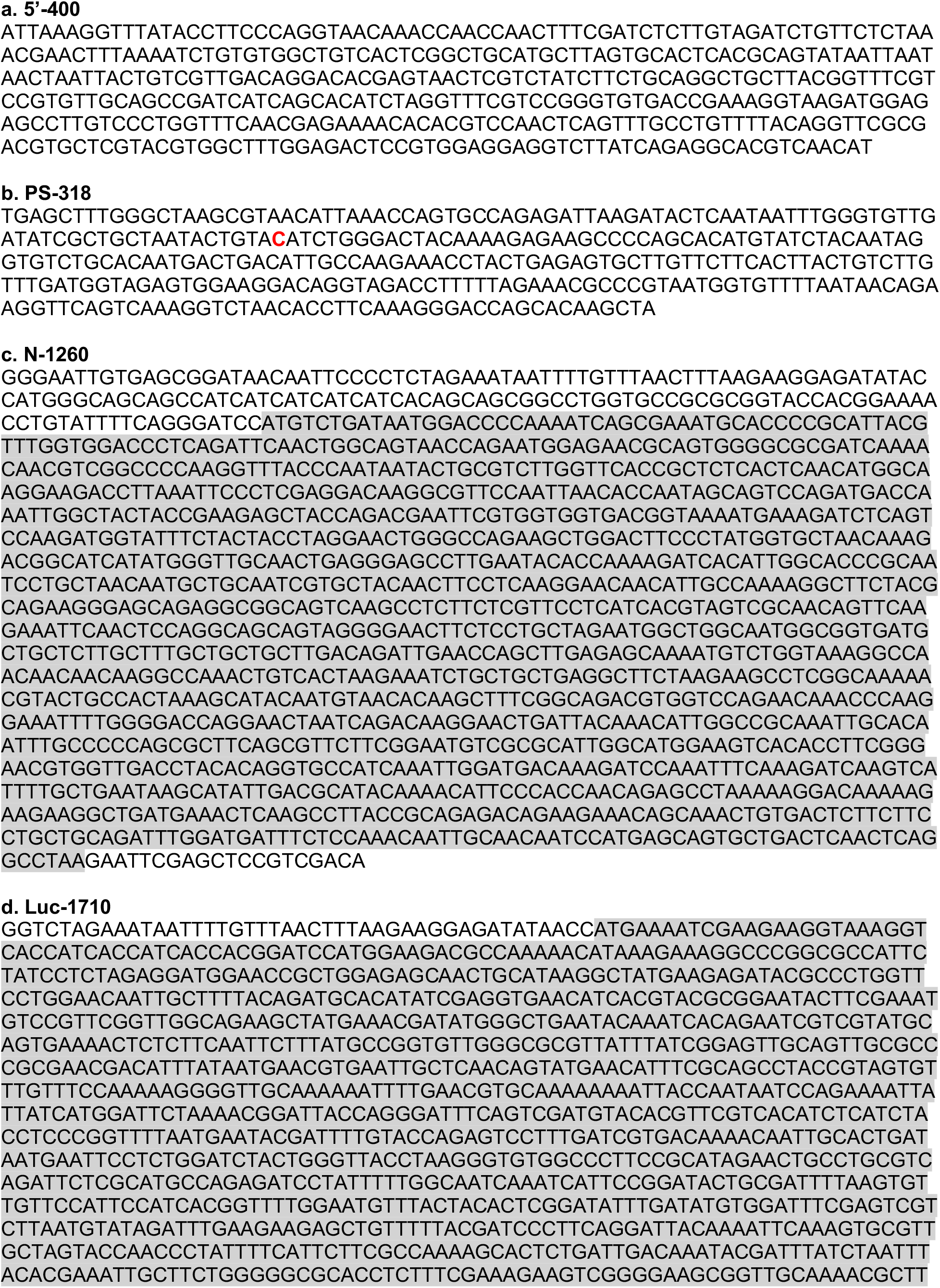
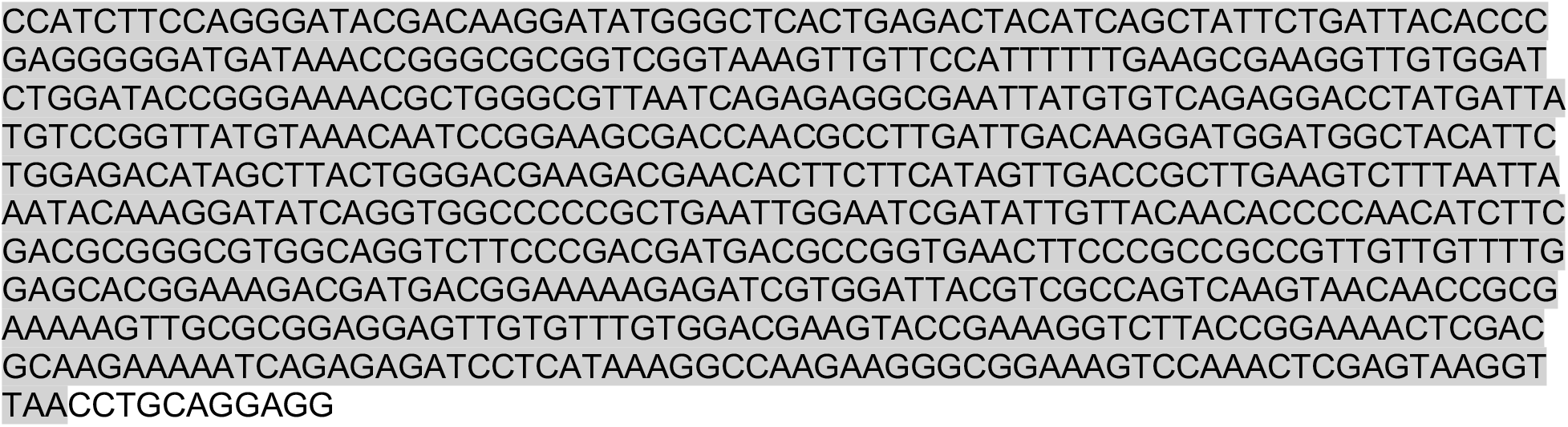
RNA sequences used in this study. **a,** 5’-400 RNA from SARS-CoV-2 (Wuhan Hu-1 strain; nt 1-400). **b,** PS-318 RNA from SARS CoV (Tor2 strain; nt 19715-20031), with an extra C (red) after A19802 as in Woo et al^55^. **c,** N-1260 RNA containing the open reading frame (gray highlight) of the N gene from SARS-CoV-2 (Wuhan Hu-1 strain; nt 28274-29533), plus flanking plasmid sequence. **d,** Luc-1710 RNA containing the firefly luciferase open reading frame (gray highlight) plus flanking plasmid sequence.

## Notes

### Competing Interest Statement

The authors have declared no competing interest.

